# Entropy Regularized Deconvolution of Cellular Cryo-Transmission Electron Tomograms

**DOI:** 10.1101/2021.04.26.441469

**Authors:** Matthew Croxford, Michael Elbaum, Muthuvel Arigovindan, Zvi Kam, David Agard, Elizabeth Villa, John Sedat

## Abstract

Cryo-electron tomography (cryo-ET) allows for the high resolution visualization of biological macromolecules. However, the technique is limited by a low signal-to-noise ratio (SNR) and variance in contrast at different frequencies, as well as reduced Z resolution. Here, we applied entropy regularized deconvolution (ER DC) to cryo-electron tomography data generated from transmission electron microscopy (TEM) and reconstructed using weighted back projection (WBP). We applied DC to several *in situ* cryo-ET data sets, and assess the results by Fourier analysis and subtomogram analysis (STA).

---

Recent advances in cryo-electron tomography (cryo-ET), most notably the ability to thin cryo-preserved specimens using a focused ion beam (FIB), have opened windows for the direct visualization of the cell interior at nanometer-scale resolution (1–9). Cells are rapidly frozen to achieve a vitreous form of ice that preserves biological molecules in a near-native state. They are then cryo-FIB milled to a suitable thickness of 100-350 nm for imaging with transmission electron microscopy (TEM). A series of projection images is acquired, typically with 1-5 degree increments and then reconstructed into a 3D volume (10). This 3D reconstruction is rendered for display and analysis, which may entail segmentation to highlight extended structures or averaging of sub-volumes for enhancement of molecular-scale resolution (11– 13).

While cryo-ET offers unparalleled resolution of cellular interiors, it is challenging for a number of reasons. First, vitrified biological samples are highly sensitive to damage by the electron irradiation required for imaging. Constraints on the permissible exposure result in limited contrast and a low signal to noise ratio (14). Additionally, higher resolution information is degraded by radiation damage over the course of imaging (15), though approaches such as dose-symmetric acquisition have been developed to optimize recording of high frequencies (16). Second, the modality of wide-field TEM depends on defocus to generate useful phase contrast, but with a non-trivial dependence on spatial frequency that is expressed in a contrast transfer function (CTF). Contrast is lost at low spatial frequencies and oscillates at high spatial frequencies, meaning that material density could be represented as intensity either darker or lighter than background (17–19).

Post-processing is applied to correct this representation in the image intensities. The correction is inherently approximate, and is especially challenging in tomography where the defocus varies across the field of view for tilted specimens (20). Third, the available raw data are never sufficient to produce an unambiguous reconstruction. The tilt range is restricted by the slab geometry, typically to about 120° around the vertical. The projected thickness of a slab also increases with tilt angle, resulting in degraded contrast and resolution from these contributions to the reconstruction. The missing information is best recognized in Fourier space, where it is known as the missing wedge. The gaps between discrete tilt angles also leave small missing wedges as seen in Fig. 1. Since the reconstruction is equivalent to an inversion in Fourier space, it is obvious that some interpolation is required and that the data are incomplete. As such, it is not surprising that different algorithms can generate somewhat different reconstructions from the same data. Commonly recognized artifacts are elongation along the Z direction and streaks projecting from high contrast points into neighboring planes in the volume. In addition to the missing wedges, TEM images require a significant defocus to get adequate contrast. For *in situ* cryo-ET data, a typical defocus of at least 5 *μ*m is used. Finally, the process of reconstruction by weighted back projection (WBP) introduces well-known problems. These include significant intensity above and below the sample volume, where we expect vacuum with no signal. This is due to cross-terms in the WBP coming from the tilt wedges, as well as distortions in the WBP arising from the missing wedge. Because of these issues with cryo-ET data, filters to improve contrast and compensate for the missing wedge are an area of ongoing research (21). These techniques include non-linear anisotropic diffusion (NAD), convolutional neural networks based on detector noise models, wavelet based filtering methods, different implementations of deconvolution, and model based iterative reconstruction (MBIR) (22–31). Here, we present a deconvolution approach to achieve both enhanced SNR and missing wedge compensation.

**Fig. 1.**
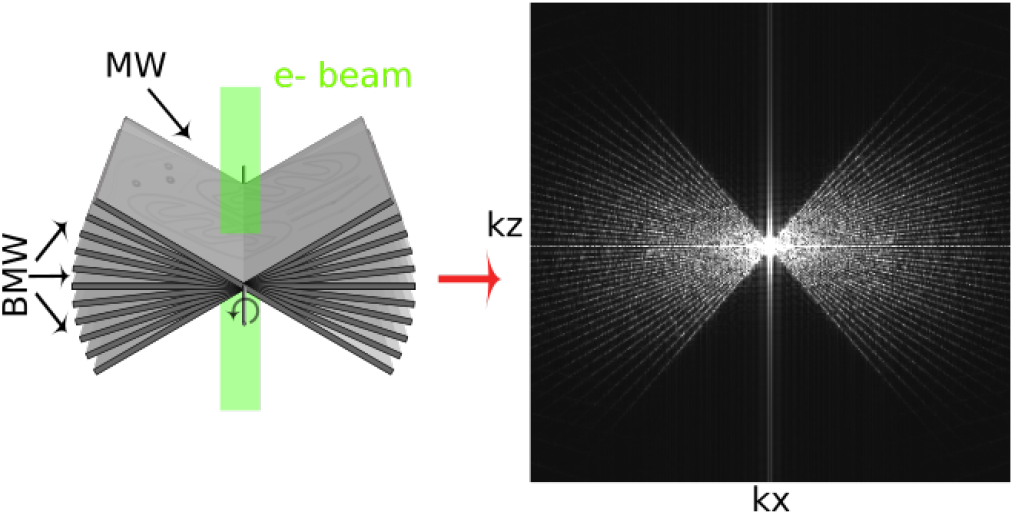
Tilt series collection and the missing wedge issue. Left: Schematic of tilt-series collection scheme. Sample projections are acquired over a range of tilt angles, typically from -60° to +60°. Right: Middle slice of the kxkz plane shows the missing wedge (MW) and baby missing wedges (BMW) of information visualized in Fourier space. Attenuation of high frequencies due to radiation damage as tilt series progresses not depicted (see (15, 16)).

The image distortions resulting from the incomplete tilt series and CTF can be characterized in terms of a single sample point in the data. This model is referred to as the point spread function (PSF), of which the hour-glass PSF in light microscopy is a classical example (32–34). Formally, the PSF is convolved with all points in the specimen function to form what is recorded in the image (35). If the PSF is well defined, it becomes possible to partially reverse the process of convolution to obtain an improved reconstruction. This reversal is referred to as deconvolution, which is a mathematical/computational iterative inversion processing procedure, extensively utilized in astronomy, spectroscopy, and light microscopy to partially restore data distorted by the imaging process (35). The deconvolution process is constrained. The most common constraint is the imposition of positivity of the deconvolved data (35). Other stabilizing constraints may include smoothing in real space to suppress high-frequency oscillations. Deconvolution is also very sensitive to noise, and most deconvolution algorithms include regularization parameters whose values are difficult to evaluate theoretically. Additionally, in most cases the deconvolution algorithms will diverge with increasing iterations, building up mottle and noise that obscure the interpretation of the final deconvolved image. Finally, most deconvolution implementations do not have a practical estimate of the error in the converged solution.

Entropy-regularized deconvolution (ER-DC) (36) is formulated to handle data with a weak signal-to-noise ratio, with a regularization term that exploits certain characteristics specific to images originating from crowded molecular environments such as cells. Specifically, in cellular images, high intensities and high second-order derivatives exhibit certain sparse distribution, and this property is exploited by the custom regularization used in ER-DC (36). This regularization was originally designed for fluorescence images, and this approach was taken recently for processing of STEM cryotomography (CSTET) reconstructions (29).

Since TEM is currently the dominant modality for biological 3-D imaging of cells (37) and its CTF is complex, it deserves a separate study, which is the focus of this paper. The major distinction is that the contrast inversions, which were absent in the STEM data as acquired for tomography, should be accommodated in the construction of the 3D PSF for TEM tomography. Here, a similar approach is taken for TEM. The combined effect of the 2D CTF of the system and the missing wedge are captured in the form a of 3D PSF. We attempt to remove the distortions caused by the 2D CTF and the missing wedge by deconvolving the back-projected 3D image with the aforementioned 3D PSF. We apply this approach to both real data, shown below, as well as to idealized simulated data, discussed in the supplement (SI Appendix Fig. 6,7) and examine the effects of ER-DC on in real space qualitatively, by Fourier analysis, and by subtomogram averaging.

## Results

### Electron Tomography Point Spread Function

The key to a meaningful deconvolution is that the synthetic PSF should represent as closely as possible the 3D image of an ideal point source. In the case of TEM, this requires an accounting for the defocus imposed in the image acquisition, which is customarily expressed in terms of a contrast transfer function (CTF). The 3D PSF for deconvolution was computed from simulated projections of a point source with the same dimensions and pixel spacing as the aligned tilt series (Fig. 2A). The CTF was first convolved with a projected point-source (Fig. 2B), and then a synthetic tilt series was reconstructed to the same dimensions as the original tomogram using the tilt angles represented in the corresponding reconstruction (Fig. 2C). This is the real-space PSF, whose 3D FFT serves as the optical transfer function, or kernel, for the deconvolution (Fig 2D). The 2D CTFs vary with the gradient of defocus for each micrograph in the tilt series.

**Fig. 2.**
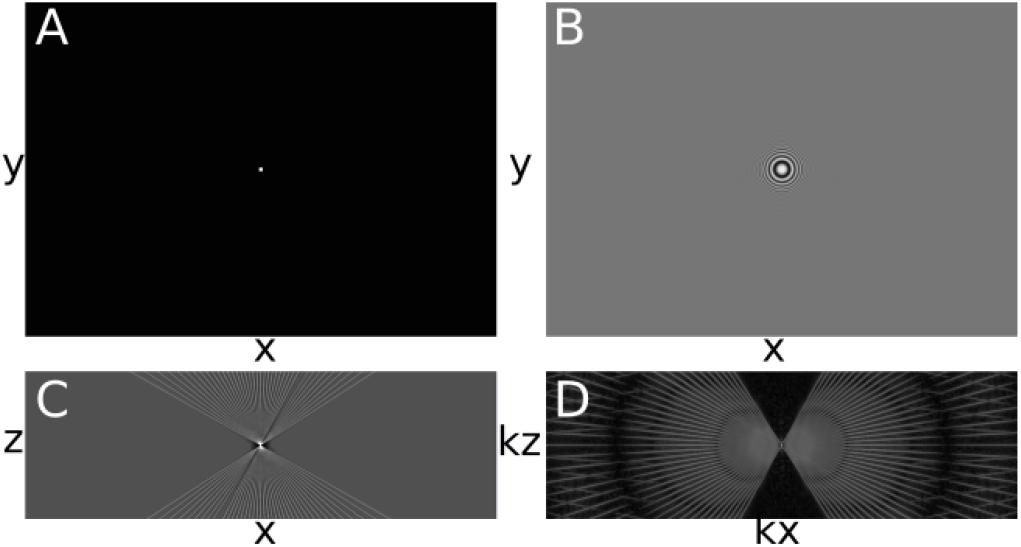
Generating the TEM Point Spread Function. (A) Synthetic tilt series of a centered point source. (B) Point source tilt series convolved with CTF. (C) Slice of the weighted WBP tomogram of convolved CTF-point source (PSF), xz view. (D) 3D FFT of tomogram showed in C, xz slice.

### Tomogram Deconvolution

As a first demonstration of cryo-ET deconvolution we used a HEK cell cultured on-grid that had been FIB-milled to 150 nm thickness. The reconstructed volume contains membranes, microtubules, and a prominent crystalline protein array. The cells were overexpressing human Parkinson’s related protein LRRK2-I2020T (38), and the observed repetitive structure is likely an autophagosome, given its double lipid bilayer structure (39). Contrast is sharp in slices through the XY plane of the tomogram, as expected (blue plane-mid structure, Fig. 3B), but contrast and resolution in the Z direction, seen in a slice through the XZ plane (orthogonal green plane in mid structure, Fig. 3C) are severely compromised. Furthermore, the reconstructed volume displays a signal both above and below the specimen when observed in the XZ plane. Since the milled slab of material is finite in the z direction, and the sample is imaged in a vacuum, there should be negligible intensity outside the sample volume in the reconstructed data. This is a known artifact of WBP. These image distortions in real space can also be characterized in Fourier space, where the real space dimensions (*x, y, z*) correspond to the Fourier dimensions (*K*_*x*_, *K*_*y*_, *K*_*z*_). The protein array in the real space XY plane appears as a lattice of calculated diffraction spots in the plane (*K*_*x*_, *K*_*y*_), as expected (Fig. 3E). In the XZ plane, the lattice of spots is sharply truncated at the Fourier planes normal to the limits of the acquired tilts. In summary, WBP suffers from major distortions visible in both real and Fourier space.

**Fig. 3.**
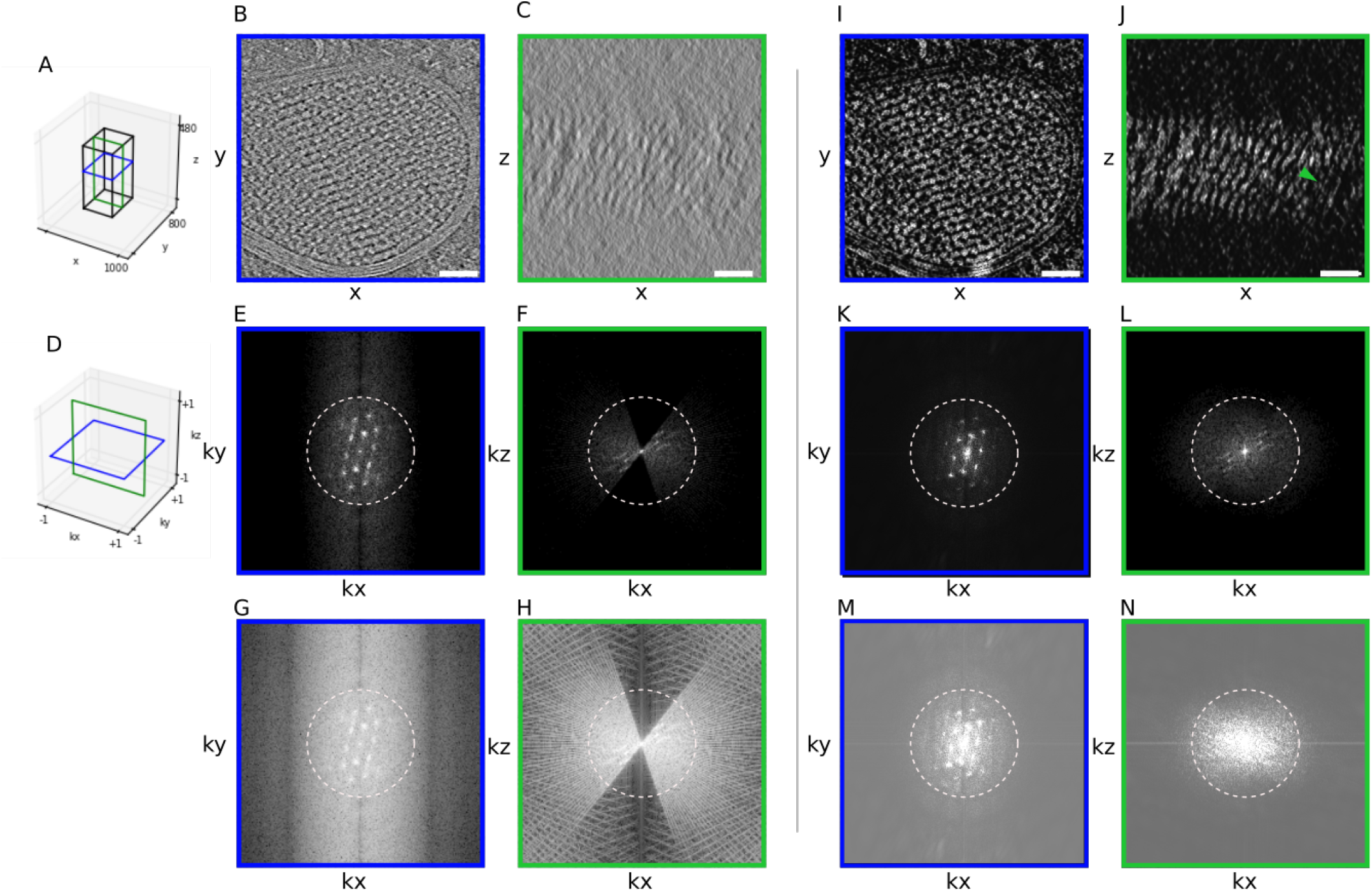
Filling of Missing Wedge by Deconvolution. (A) Schematic of slices used to generate panels B, C, I and J. (B)XY Slice of a tomogram of a HEK cell reconstructed using WBP. Throughout this work, white intensities correspond to high density values. (C) XZ slice of the same tomogram. (D) Schematic to show slices through Fourier space used to generate E,G,K,M in blue and F,H,M,L,M in green. (E,G) Slice through 3D FFT of the WBP corresponding to XY shown with a different distribution of voxel intensities. (F,H) Slice through 3D FFT of the WBP corresponding to XZ shown at two different intensity scales. (I-N) Corresponding results for the tomogram after deconvolution. Scale bars: 100 nm. For reference, dashed circles indicate 0.03 Å^*−*1^ resolution.

The result of 3D deconvolution is shown alongside the reconstruction in Fig. 3. Full details appear in the Supporting Information. All processing was performed using the PRIISM image processing software (40). Briefly, the entropy-regularized deconvolution algorithm from PRIISM was applied using the simulated PSF. While contrast is enhanced in the XY plane, the more striking improvement is seen in the XZ plane (Fig. 3J) in comparison with the WBP (Fig. 3C). In the deconvolved tomogram, two lipid bilayers are visible (arrow) across the entire sample along Z, as is the crystalline array (Fig. 3J). The restoration of information along Z in real space can also be seen in the 3D Fourier transform of the deconvolved volume, which shows increased signal in the previously empty regions corresponding to the missing wedges (Fig. 3F,L).

A very effective way to observe the results of deconvolution is to study a small volume of the WBP and/or deconvolved in a dynamic interacting display module, typically a video of the rotating volume. Stereo pairs with additional rotated views are shown for the WBP and deconvolution (SI Appendix videos 1 and 2). These may be rocked with a cursor bar, as described in Supporting Information, in order to gain an impression in 3D. Distortions along the Z axis associated with the WBP are largely removed after deconvolution. Significant information in the power spectrum appears beyond a spatial frequency of approximately 2.5 nm, which corresponds nominally to the second zero in the CTF for a 6-*μ*m defocus.

### A Second Deconvolution Example

For a second example, we applied ER-DC to a tomogram of a relatively thick lamella of *S. cerevisiae* cells (370 nm). Besides the thickness, cryo-electron tomography data of nuclei are challenging samples to interpret as nuclei are densely packed, and lack high-contrast features like membranes, cytoskeletal filaments, or large and defined particles such as ribosomes. As with the deconvolution of mammalian cells, deconvolution provided increased contrast in XY and an improved ability to visually interpret information along Z compared to the WBP. The nuclear envelope is clearly visible in the XY slices of the WBP and the two deconvolutions (Fig. 4 A, E, I). In XZ however, no clear structure can be followed in the BP(Fig. 4C), but can be more easily followed in the deconvolution (Fig. 4 G, K)). Additionally, the missing wedge seen in Fourier space is filled in by the deconvolution process (Fig. 4 H, L)). By utilizing rotating angle stereo-pair renderings of the volume (RAPSs), one can compare the WBP and deconvolved volumes in 3D (SI Appendix videos 3 and 4). In the BP, there is little distinguishable structure as the volume rotates. In contrast, fine features can be identified at every angle, such as the nuclear envelope, as well as densities that could correspond to chromatin and nucleosomes. The 3D-FFT of the deconvolution (Fig. 4), KxKz view, shows the missing wedges being filled in, indicating that the deconvolution process helps correct for these artifacts, even in challenging samples.

**Fig. 4.**
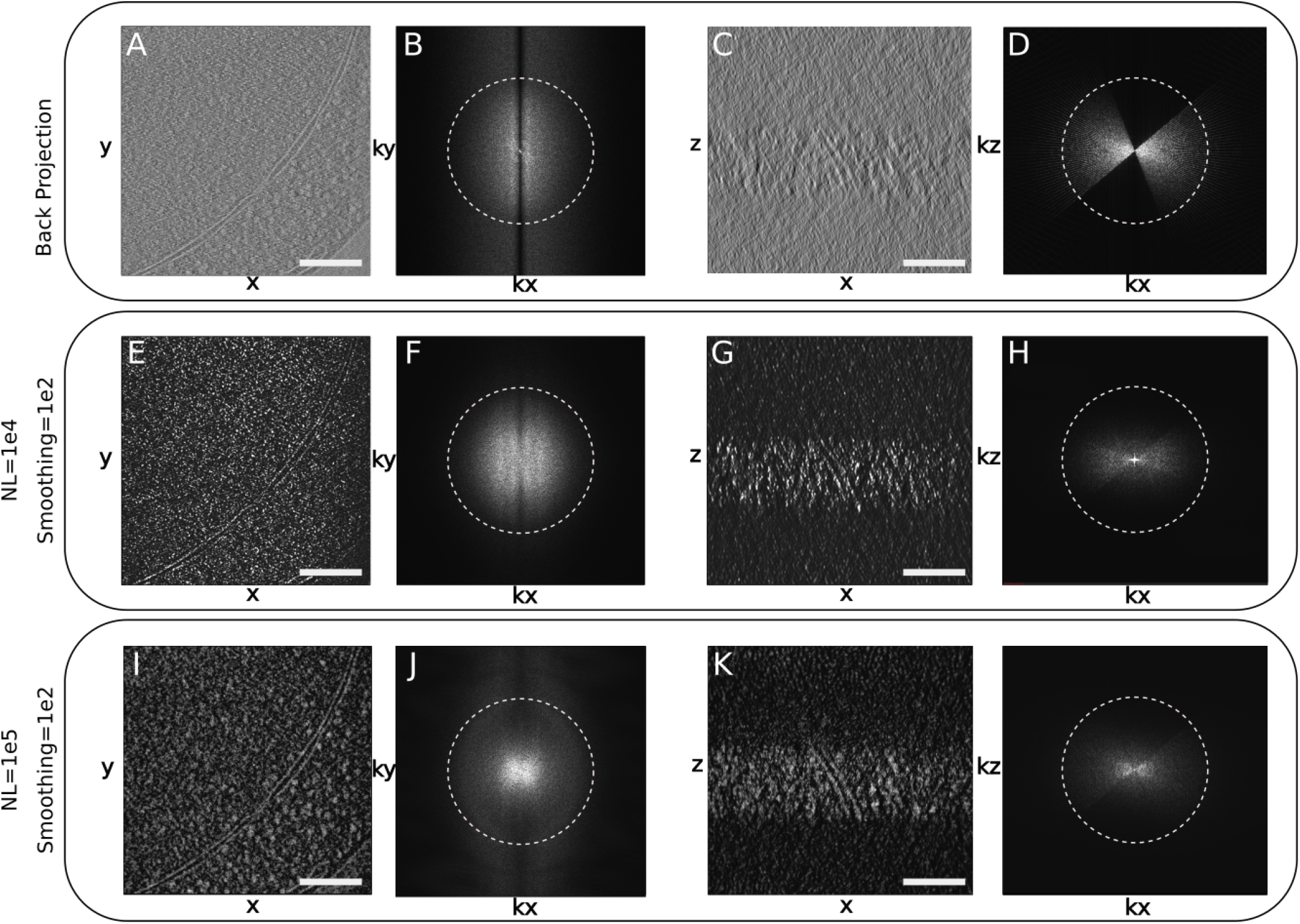
Deconvolution of a Tomogram of the Nuclear Periphery of a *S. cerevisiae* Yeast Cell. (A) Central slice (10.6 nm thick) of the XY plane of a WBP tomogram. (B) Fourier transform of A. (C) Central slice of the XZ plane of the WBP tomogram. (D) Fourier transform of C. (E) Central slice of the XY plane of the tomogram deconvolved with a smoothing parameter of 100 and a non-linearity factor of 10,000. (F)Fourier transform of E. (G) Central slice of the XZ plane of the deconvolved tomogram. (H) Fourier transform of G.(I) 10.6 nm slice of the XY plane of a the tomogram from A, deconvolved with a smoothing parameter of 100 and a non-linearity factor of 100,000. (J)Fourier transform of I. (K) 10.6 nm slice of the XZ plane of the deconvolved tomogram from I. (L) Fourier transform of K. Scale bars: 100 nm, dashed circles indicate 0.03 Å^*−*1^ resolution.

### Deconvolution and Subtomogram Analysis

Subtomogram analysis is an approach to protein structure determination *in situ* (11, 41–55). Similarly to single particle analysis, of which it is an extension to 3D, averaging multiple examples of images that represent particles of the same kind serves to reduce noise. If the molecules lie in random orientations, 3D averaging can also be used to compensate for the missing wedge (11). The crystalline-like body seen in Fig. 3 provided an interesting test case for averaging where orientations were determined to be uniform by translational symmetry in the crystal (Fig. 5). Therefore, only select orientations are represented in the sample. First, we attempted to align the crystal subunits over 360^0^ in *θ* and *ϕ* on the WBP reconstruction. This resulted in an alignment that was dominated by the missing wedge, a common pitfall in subtomogram averaging, and produced a structure that was strongly elongated(Fig. 5A). Second, we used the same particles, but this time from the deconvolved data set. Alignment and averaging resulted in a structure that resembled much better the unit of the crystalline array in the original tomogram (Fig. 5E). Third, we averaged the WBP particles using the transformations determined by the deconvolution alignment. In this last case, we obtained a structure similar to the one obtained from deconvolution-aligned and deconvolution-averaged particles (SI Appendix Fig. 4B), demonstrating that the alignment of subtomograms is improved by deconvolution. This tomogram was acquired from a HEK cell overexpressing human LRRK2 (38). While the identity of the molecules forming the crystalline-like array was not specifically established (*e*.*g*., by CLEM) and the number of particles in this tomogram is severely limited (82), the overall shape of the deconvolved average resembles the cryo-EM structures of LRRK2 determined both *in situ* bound to microtubules (38) and *in vitro* (56).

**Fig. 5.**
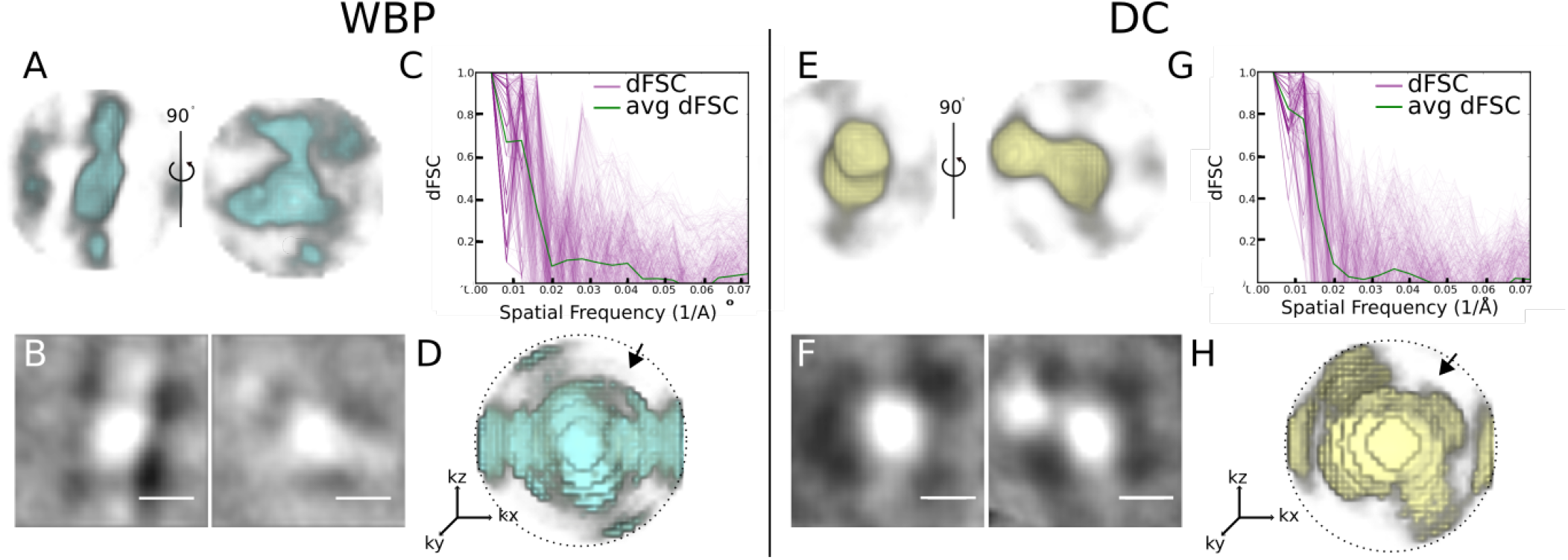
Comparison of subtomogram analysis of a crystalline body in WBP vs deconvolved reconstructions. (A) Two views of the crystal body average of WBP subtomograms. (B) Central 1.4 nm slice of WBP crystal body subtomogram average. (C) 3D-FSC curves generated from two half-map averages of the WBP crystal body subtomogram average. Green line is the average, pink lines are individual directional FSCs. D) 3D render of dFSC curves; arrowhead denotes the center of the missing wedge. (E-H) corresponding averages and FSCs derived from the deconvolved volume. Scale bars: 10 nm. dFSCs shown out to Nyquist resolution (0.07Å^*−*1^).

Fourier shell correlation (FSC) is widely used in single particle cryo-EM (57), as a metric of the resolution of a molecular structure. It is a quantitative measure of similarity, typically implemented in cryo-EM by comparing two structures, each generated from a half data set. The comparison is done by correlating shells of the 3-D Fourier transform of each of the half maps. Standard FSC compares global similarities, correlating all orientations contained within a shell, giving a single curve for the entire structure as a function of spatial frequency. Resolution is then quoted as the inverse spatial frequency where the correlation drops below an accepted threshold. Directional FSC (dFSC) is a variant in which all Euler angles are explored for frequency comparison, and provides a representation of resolution in all directions (58). dFSC was applied to two half-map averages from the crystalline array in the WBP, and then in the deconvolved averages to assess changes in resolution in any direction between the WBP (Fig. 5 C) and deconvolution (Fig. 5G). The resolution from the averaged dFSC curves is similar for the deconvolved and the WBP reconstructions (∼ 53Å), using the gold-standard FSC); however, the curves of the averaged and individual dFSC have higher correlation at this and higher resolutions. This is evident when comparing the 3-D rendering of the dFSC for the WBP (Fig. 5 D) and deconvolution (Fig. 5H), where the resolution is anisotropic (lower correlation in the area of the missing wedge denoted by an arrow) for the WBP but not for the deconvolution. Further, higher resolution is found in the deconvolution reconstruction.

To investigate the effects of deconvolution in the alignment of the particles and the improvement of the average due to the missing wedge separately, we chose to use microtubules, since their structure is well established, as are the pipelines for subtomogram analysis. We analyzed a tomogram of reconstituted microtubules decorated by the Parkinson’s related protein LRRK2^RCKW^(56). In the tomogram, it is evident that the deconvolution process increased the contrast between the microtubules and the surrounding media, and we again see a reduction in XZ distortions (Fig. 6A,B,D,E), as well as a corresponding filling of information in the missing wedge in Fourier space (Fig. 6C, F). Microtubule subtomograms were extracted from both the WBP and deconvolved volumes using the filament tracing function in Dynamo(59). The subtomograms were independently aligned and averaged as described in (38) and in Materials and Methods (Fig. 7 A, B). Note that the contrast between protofilaments is distinctly sharper for the deconvolved data. However, this method of alignment includes an azimuthal randomization that is specifically designed to average out the missing wedge in the final average. To assess the effect of deconvolution specifically on the missing wedge, we ran the alignment without this randomization step, that is with the missing wedge always in the same orientation, as it exists in the original particles. Compared to the WBP, the deconvolution-processed average shows increased distinction between protofilaments in the direction of the missing wedge (Fig. 7D). Lastly, we used the alignment parameters generated from the azimuthally zeroed deconvolved subtomograms to the WBP particles to generate a WBP average (Fig. 7E). Here, the deconvolution-aligned WBP average still shows a prominent missing wedge, similar to the WBP average generated by aligning WBP particles. This indicates that the improvement in the average from the deconvolved particles is not simply due to improved alignment, but that the filling of the missing wedge is reducing distortions in XZ, thereby improving the resulting subtomogram averages.

**Fig. 6.**
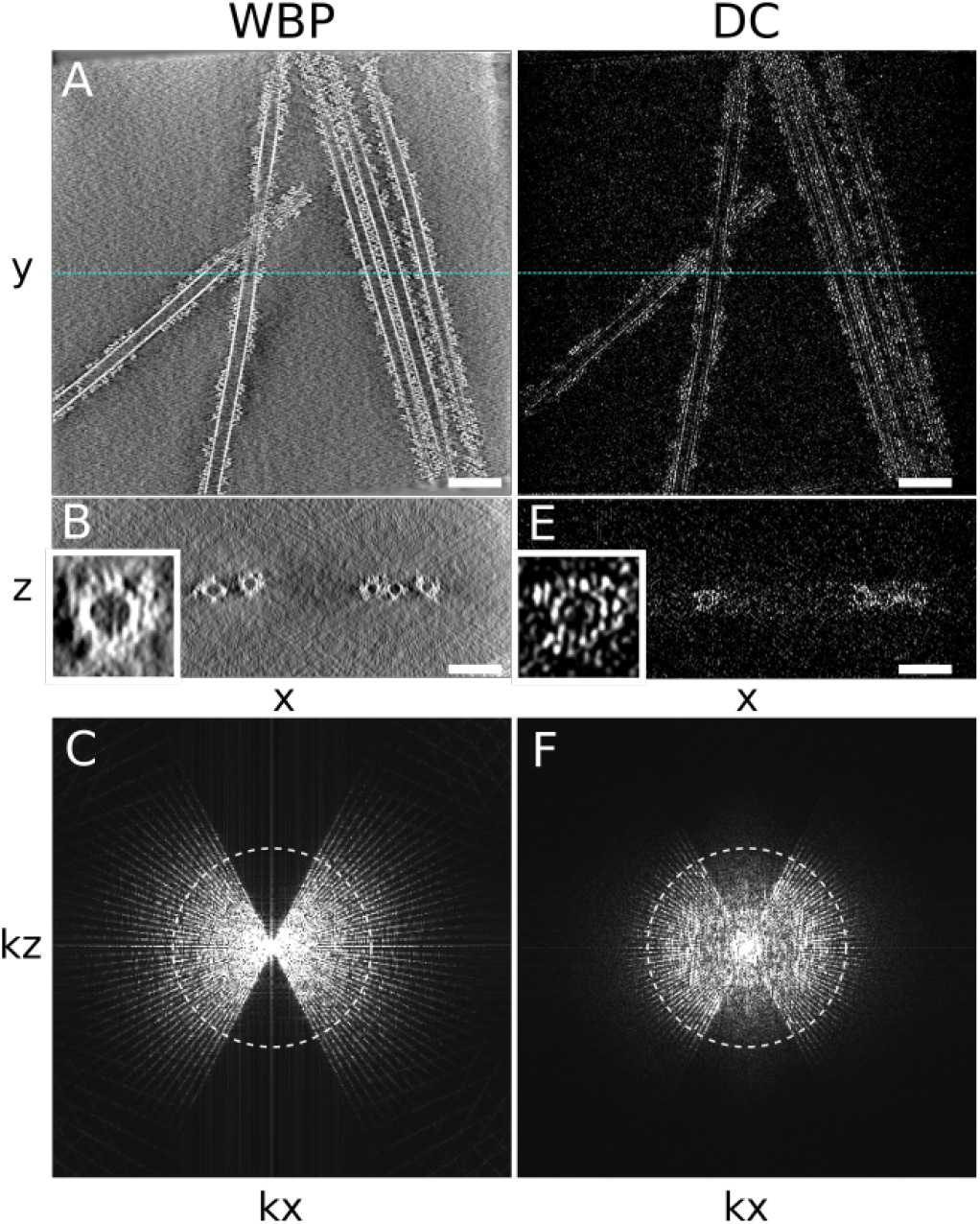
Deconvolution of a tomogram of reconstituted microtubules. (A) XY slice of the WBP-reconstructed tomogram containing microtubules. (B) XZ view of the tomogram in A; blue dashed line in A corresponds to the slice shown. (C) kxkz view showing the missing wedge. (D) XY of the deconvolution tomogram. (E) XZ of deconvolved tomogram in D; blue dashed line in D corresponds to the slice shown. (F) kxkz of the deconvolved tomogram. Scale bar = 50 nm, dashed circles indicate 0.06 Å^*−*1^ resolution

**Fig. 7.**
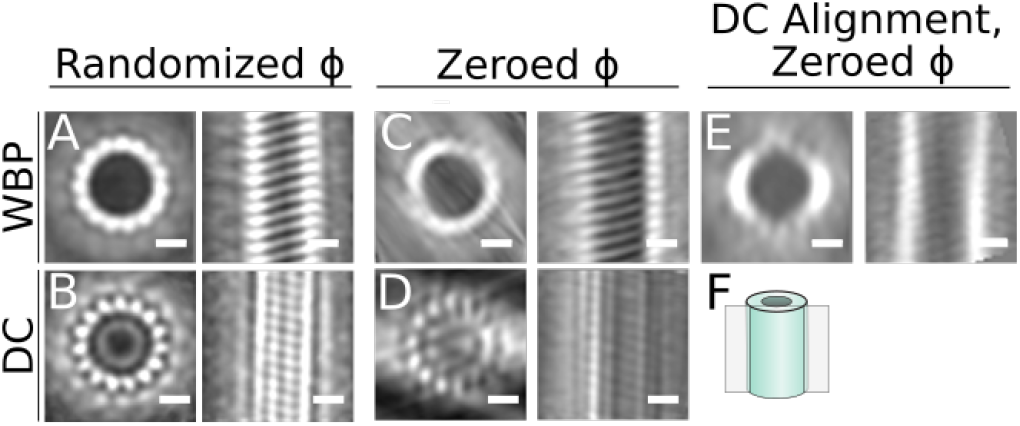
Subtomogram analysis of WBP and deconvolution-processed microtubule. In all panels, top and side views of the average are shown for the microtubule average obtained under the specified conditions (A) Average from the WBP tomogram, using a randomized azimuth-angle (*ϕ*) averaging approach to compensate for the missing wedge (38). (B) Average from the deconvolved tomogram using the same randomized *ϕ* scheme. (C) Average from the WBP tomogram, with initial constant *ϕ* angles for all particles, allowing the missing wedge to affect the average. (D) Average from the deconvolved tomogram using the same constant *ϕ* scheme. (E) Average generated by applying the alignment parameters from the deconvolution uniform starting azimuth/restricted rotation alignment to the the WBP particles. (F) Schematic showing the location of the slice on the right-hand side image in each panel. Scale bars: 10 nm.

In addition to the experimental data explored in this paper, we used simulated tomography data was to investigate the effect of DC on a system with a known solution. While the noise models in simulated data are not fully comparable to experimental data, we reasoned that we should apply DC to these data with the expectation that it recovers the initial structure accurately by filling in the missing information in the missing wedge area. We picked a X-ray crystallography structure of a microtubule from the Protein Databank (PDB-3J2U) to generate a simulated cryo-EM map at 3.3Åresolution using EMAN2 (60). Then, we simulated a tilt series of the density map (SI Appendix Fig 6), which was reconstructed using our standard WBP (SI Appendix Fig 7B). Deconvolution was applied to this simulated tomogram, resulting in a near complete refilling of the missing wedge (SI Appendix Fig 7 D,G), demonstrating that DC works in idealized cases.

## Discussion

We have successfully applied ER-DC to cryo-electron tomograms, and demonstrated enhanced contrast compared to the WBP reconstructions, as well as reduced resolution anisotropy along the Z axis. In real space, one can follow membranes in the XZ plane of the deconvolved volume that were hardly visible in the WBP. In Fourier space, it is clear that portions of the missing wedge are filled in, and the distribution of voxel intensities changes significantly as a result of deconvolution. However, there are still several considerations for TEM deconvolution, and these are further discussed in the Supporting Information.

First, the reality is that deconvolution acts as a filter for the data. The intensity of each voxel is modified in some fashion, and care must be taken in interpreting the deconvolved volume. Deconvolution has two parameters, for non-linearity and smoothness, and the optimal values must be determined experimentally by systematically varying the parameters over several orders of magnitude; the parameter search quickly settles into basic convergent deconvolved images that look biologically reasonable (e.g. membrane bilayers are visible, ribosomes are distinct, etc.). At the end of the deconvolution process, one can usually settle on a few deconvolved images coming from a close smoothness parameter. These different deconvolved images are studied side-by-side comparing 3-dimensional volumes for details. The side-by-side images are very similar to one another, but subtle features between them exist. Crucial are the orthogonal Z image planes for judging smoothness parameters and structure. There are a number of considerations for the deconvolution process, discussed in the Supporting Information.

How does one know if the deconvolved structure is credible? In addition to the side-by-side study of several smoothness deconvolved images, a control raw WBP image must be studied alongside the deconvolved images, at several intensity scalings of the WBP data. Any feature uncovered in the deconvolved data would be searched for in the raw WBP data control, and would have to be present in the WBP control. However, in our experience, the deconvolution process has never been observed to invent a structure that is not present in the WBP raw data control (29).

This study makes the statement that the missing wedge of information is substantially filled by deconvolution. Visually and in Fourier space representation this is the case; however, this statement needs caution. We do not know if deconvolution will improve Z resolution for certain kinds of data, intensities, or different structures *e*.*g*., of various sizes. It is possible that spaced periodic structures positioned on top of one another along Z in a tomogram are not resolved correctly in the deconvolved data.

A second point in the deconvolution discussion centers on what mathematics allows unobserved data to propagate from areas of observed data, into their correct structural space. Since all image information can be decomposed into 3-dimensional Fourier representation, one is, in essence, saying that there is information in one region of Fourier space that can be extrapolated correctly into other regions of Fourier space by the deconvolution process. There are two examples from the inverse problems literature to reassure that such extrapolated information can indeed be real. The first one is called “Analytical Continuation” ((61), and references therein), which is known in optics literature. The Analytical Continuation (AC) conjecture, taken from (61), states: All image information can be decomposed into a Fourier transform, and a spatially bounded region of Fourier space can be expressed as an analytical function. The analytical function can be exactly known for a small region, and if there is no noise, the entire analytical function can be determined/extrapolated by AC. The extension can continue indefinitely, and this is a hallmark of AC (61). Noise is critical and the analytical values become small as iterations progress as the function get extended to higher resolution regions of Fourier space, reasons AC is little used (but see (61)). In the case of ER-DC, noise is heavily suppressed and resolution extensions required are modest, suggesting that AC might work.

The second one is called compressive sensing reconstruction, used in modalities such as magnetic resonance imaging (62) and tomography (63). It involves high-quality reconstruction from highly under-sampled Fourier data and tomographic projections with a limited set of angles with regularization constructed using derivatives. Because of the way the derivative operator is related to the measurement operator (tomographic projection or Fourier transformation), high quality reconstruction becomes possible from sparse Fourier samples or from tomographic projections from a limited set of angles. Although these theories are not directly extensible to our recovery problem, they reassure that extensions in Fourier space are possible, and hence the filled-in missing wedges may be trusted if the resultant structures in the real space appear plausible.

Another independent argument supporting why the missing wedges could correctly be filled can be given from a statistical viewpoint. Recall that the regularization used in the ER-DC enforces certain hypothesized joint distribution of intensity and second-order derivative magnitude. It turns out that the back projected images deviate significantly from this joint distribution. Hence, the minimization involved in ER-DC brings in a proper filling on the wedges such that: 1) the resulting real space image is consistent with the measured projections, and 2) the resulting real space image better matches with hypothesized distribution.

Thirdly, deconvolution could have an impact on the electron dose required to obtain a suitable tomogram. The deconvolution process might allow other dose reduction steps, such as fewer tilts and lower beam intensity. In addition, there are several aspects of the deconvolution process that can be improved, and are described in the Supporting Information. The deconvolution process filling in the missing wedges in Fourier space allows biological structures to be followed in 3-dimensions. This resolution is adequate to see, for example, gaps between the 10 nm nucleosomes allowing a chromosome path to be followed. One imagines a two step process for cellular tomography: first, the path of a structure is followed with the architecture discerned, a process greatly improved by deconvolution. Subsequently, once an architecture is determined, molecular features can be superimposed using averaging methods and molecular modelling (38).

## Materials and Methods

### Sample Preparation

Yeast *S. cerevisiae* W303a cells were grown at 30°C in YPD media (1% yeast extract, 2% bactopeptone, and 2% glucose) to mid-log phase, after which 5-*μ*L were deposited in a glow-discharged Quantifoil grid (200-mesh copper R2/1, Electron Microscopy Sciences), followed by manual blotting and plunge freezing in a 50/50 ethane propane mix (Airgas) using a custom-built manual plunger (Max Planck Institute of Biochemistry). Human Embryonic (HEK-293T) cells transfected with LRRK2-I2020T cells were prepared as described in (38). *In vitro* reconstituted LRRK2-I2020T was prepared as described in (56). For both yeast and HEK cells, frozen cells were micromachined on a Scios or an Aquilos 2 DualBeam FIB/SEM microscope (TFS). FIB milling was done as described in (6).

### Cryo-electron tomography

Tilt series were obtained on a 300 kV Tecnai G2 Polara (TFS) or Titan Krios with a field emission gun, a GIF Quantum LS energy filter (Gatan) and a K2 Summit 4k×4k pixel direct electron detector (Gatan). Tilt series were acquired between ± 50° and ± 70° with increments of 2° and 3°, total electron doses between 70 and 100 e^*−*^*/*Å^2^ at a target defocus of 5*μ*m, and a pixel size of 2.2 or 3.5Å using the SerialEM software (64) in low-dose mode. Bidirectional or dose-symmetric tomography acquisition schemes were used (65), corrected for the pretilt of the lamella where appropriate. Images acquired on the K2 detector were taken in counting mode, divided into frames of 0.075 to 0.1 s.

### Tomogram Reconstruction

Tilt series were aligned and dose-weighted by cumulative dose with MotionCorr2 (66). Dose-weighted tilt series were aligned and reconstructed using Etomo, part of the IMOD package (67). Patch tracking was used to define the model for fine alignment. The aligned tilt series were reconstructed using WBP to generate the 3D tomograms.

### Deconvolution

A set of synthetic projections was generated with x and y dimensions and pixel spacing matching the tilt series that was used to make the original reconstructed volume. Each projection had a centered point source that is then convolved with the inverse Fourier transform of the CTF, generated using the defocus and astigmatism parameters estimated by CTFFIND4 (20). The convolved point source/CTF is then reconstructed using the same WBP used to generate the target reconstructed volume. Finally, the reconstruction is cropped to the same dimensions as the volume to be deconvolved, the 3D FFT of which will be used as the final PSF. Deconvolution is then run for 100 cycles using the generated PSF. A detailed description of the deconvolution procedure can be found in the Supporting Information.

### Subtomogram Analysis

Microtubule filaments were traced in Dynamo (59) to define coordinates and orientation. Single particles were defined every 4 nm along the filament, and subtomograms with a side length of 66 nm were then extracted from both the back projected and the deconvolved tomograms using these coordinates. For both sets of particles, subtomograms were iteratively aligned over three rounds of two iterations each. The particles were aligned using a spherical alignment mask to minimize bias. For the first round, the alignment was constrained to a 180 degree cone aperture, with no flip allowed and 20 degrees of azimuthal rotation, corresponding to the third Euler angle. Rounds two and three used a 30 and 10 degree cone aperture, respectively, and an azimuthal search range of 10 and 2 degrees respectively. No symmetry was assumed in the alignment. For further details, see (38). To assess any compensation for the missing wedge, alignment was performed on particles with initial tables describing the particles orientation from 1) a blank table to set all particle orientations to zero, and 2) a random table assigning each particle a random orientation.

To calculate averages for the autophagosome crystal subunit in the WBP and deconvolution tomograms, first 50 particles were identified manually in the deconvolved volume to generate an initial average. This initial average was used as a template for Dynamo’s template matching functionality and used to search for similar particles. A cross correlation threshold of 0.38 was selected, below which many particles appeared as false positives by visual inspection. Using the coordinates and putative orientations from template matching, 82 particles were cropped from both the back projected and deconvolved volumes. A global alignment was used on each data set in two (even and odd sets) using the Dynamo subtomogram alignment function (59). Each half data set was averaged and the directional Fourier shell correlation (dFSC) between the resulting half-averages(58). The alignment angles from the deconvolved particles were then applied to the WBP particles to create the average shown in Fig. 5A and to the relative resolution by dFSC.

### Simulated data

To validate the effect of deconvolution on the missing wedge under ideal circumstances, a microtubule tomogram was generated from an existing crystal structure (PDB 3J2U). After removing the chains that did not correspond to tubulin subunits forming the microtubule from the model in UCSF Chimera (68), the PDB map was converted to an electron density map using the EMAN2 functionality *e*2*pdb*2*mrc*.*py* to convert it to a simulated electron density map, followed by the *e*2*spt*_*simulation*.*py* function to simulate a tilt series (60). At this stage, the simulated tilt series was an idealized example, with no CTF applied. *e*2*spt*_*simulation*.*py* defaults to simulating the particle as if it were embedded in 400 nm of vitreous ice, and the tilt series was binned to make the pixel size 3.3 Å / px to approximate the sampling often used in cellular tomograms. The CTF was simulated and applied to the synthetic tilt series with Priism’s *pf ocusramp*. The CTF parameters included a defocus of -3.00 um, with no astigmatism. A corresponding point spread function was generated by applying the same CTF to a simulated point source tomogram derived from the simulated microtubule tilt series. The simulated tilt series was reconstructed by weighted back projection, then deconvolved with the corresponding PSF.

## Data Availability

The tomograms and their corresponding deconvolutions have been deposited in the Electron Microscopy Database. Yeast WBP and deconvolution data can be found at EMD-24433, EMD-24434 respectively. The tilt series and the corresponding tilt and defocus files are deposited at EMPIAR-10762. The inclusion body WBP and deconvolution data can be found at EMD-24435, EMD-24436 respectively. The tilt series and the corresponding tilt and defocus files are deposited at EMPIAR-10761. All the wrapper scripts necessary to perform the steps described here available at https://github.com/Villa-Lab/ER-DC.

## Acknowledgements

We are most indebted to Eric Branlund whose computational efforts made this work possible. We used samples and tomograms provided by Robert Buschauer, Reika Watanabe, Colin Deniston and Andres Leschziner. We thank Drs. Yifan Cheng (UCSF), David DeRosier (Brandeis & La Jolla), Robert Stroud (UCSF), Susan Taylor (UC San Diego), and members of the Villa lab for discussions and support. ER-DC software and documentation can be obtained from Agard/Sedat at UCSF. This work was supported by an NIH New Innovator Award (DP2 GM123494) and an NSF MRI grant (DBI 1920374) to E.V., an NIH R35 grant to D.A.A. (NIH R35GM118099) and private funds from J.S. ME is the Sam and Ayala Zacks Professorial Chair and head of the Irving and Cherna Moskowitz Center for Nano and Bio-Nano Imaging. We acknowledge the use of the UC San Diego cryoEM facility, which was built and equipped with funds from UC San Diego and an initial gift from Agouron Institute, and of the San Diego Nanotechnology Infrastructure (SDNI) of UC San Diego, a member of the National Nanotechnology Coordinated Infrastructure, supported by the NSF grant ECCS-1542148.

## Notes

### Competing Interest Statement

The authors have declared no competing interest.

