## Supplemental Information for "Entropy Regularized Deconvolution of Cellular Cryo-Transmission Electron Tomograms"

#### This PDF file includes:

Supplementary text
Figs. S1 to S7
Legends for Movies S1 to S4
SI References

#### Other supplementary materials for this manuscript include the following:

Movies S1 to S4

### Supporting Information Text

### Detailed Deconvolution Workflow

**Computing the Transfer Function for the Deconvolution.** To include the microscope contrast transfer function (CTF) in the transfer function for deconvolution, the per-projection defocus and astigmatism parameters were determined by running CTFFind 4.0.7 (1) for each of the projections in the aligned and motion-corrected tilt series. The parameters used for CTFFind were the tomogram pixel spacing (2.9 or 3.6 Å, depending on magnification), 300 kV for the acceleration voltage, 2.27 mm and 2.7 mm for the spherical aberration of the Polara and the Titan $K_r$ios respectively, 0.07 for the amplitude contrast, 512 for the size of the power spectrum, 44 and 11 Å for the minimum and maximum resolutions respectively, 10000 and 100000 Å for the minimum and maximum defocus respectively, 2500 Å for the defocus search step, and 100 for the expected astigmatism. For those projections whose radially averaged spectra from CTFFind did not have clear rings from the zeros in the contrast transfer function, the means of the defocus and astigmatism parameters from the other projections were used.

The transfer function for deconvolution were computed with these steps (SI Appendix Fig. 1):

1. Generate synthetic projection data for a centered point object interacting with the electron beam. That synthetic projection data incorporated the per-projection defocus estimates from CTFFind.

2. Reconstruct that synthetic projection data with the same reconstruction method as used for the experimental tilt series.

3. Compute the three-dimensional discrete Fourier transform of the result of step two to get the unnormalized transfer function.

4. Divide the values in the unnormalized transfer function by its value at the zero frequency so that applying the transfer function to a sampled volume does not change the volume's mean intensity value.

The result of step two is the sampled point spread function (PSF) for the combined process of imaging the point object at different tilt angles and reconstructing it.

Synthetic projection data were generated in two stages. First, a series of images are generated to match the number of projections and the dimensions of the measured tilt series and the dimensions of the tilt series after binning by two and trimming for the reconstruction region of interest. All samples in those images are zero except for the central pixel which has a value of one. Those images are then blurred by the modeled microscope CTF incorporating the per-projection defocus estimates. The initial point source tilt series is made using PRIISM 4.6.1, and a text file, `tilts.txt`, that has the tilt angles, one per line and in units of degrees, for the measured tilt series. This was achieved by first taking the 2D FFT of each projection of the aligned tilt series, then thresholding the result to give a single pixel central point with an intensity value of 1.

```
insert_tilts tiltseries.ali tilts.txt
FTransform2D tiltseries.ali tiltseriesfft.mrc -center_zero -real_complex_full -same_units \
    -x=0:(Xdim-1) -y=0:(Ydim-1)
Threshold tiltseriesfft.mrc synproj_noblur.mrc -result=mask -not_below=6 -mode=short
```

Using the file named `synproj_noblur.mrc` from the first stage, `pfocusramp` from PRIISM 4.6.1, and a text file named `defocus.txt` with the estimated defocus values, one per line in units of microns and written with a negative sign to match PRIISM's convention for defocus, the second stage can be implemented by running `pfocusramp` as follows:

```
pfocusramp synproj_noblur.mrc synproj_blur.mrc -amp=0.07 -axis=0 -cs=2.27
    -deftxt=defocus.txt -kv=300 -mtf=1:5.2279:0:50 -op=apply
```

That includes a Gaussian envelope, $e^{-5.2279q^2}$ where q is the spatial frequency in cycles per sample, to approximate the observed loss of power in the radially averaged spectra from CTFFind.

The weighted back projection of the blurred synthetic projection series was computed to get the PSF using PRIISM 4.6.1's `ewbp` as follows:

```
ewbp synproj_blur.mrc psf.mrc -reconxz=1000:480 -sizexz=1000:480 -iy=0:799
    -filter=2 -hdfilt=1 -moderec=2
```

The command for the measured tilt series specified the sizes in terms of the unbinned coordinates; here they are in the binned coordinates and are one half the size.

PRIISM 4.6.1's FTransform3D was used to compute the three dimensional discrete Fourier transform of the PSF to get the transfer function:

```
FTransform3D psf.mrc tf.mrc -same_units -shift=499:240:399
```

In computing the Fourier transform, the origin of the spatial domain was shifted to match the peak of the PSF so that applying the transfer function to a volume does not shift the contents of the volume. Also, to match what is expected by the deconvolution software, the metadata for the sample spacing was kept in the spatial domain units rather than converting it to the equivalent spacing in the frequency domain.

**Computing the Weighted Backprojection to be Deconvolved.** The tomogram to be deconvolved was reconstructed by elliptically weighted back projection from the aligned and motion-corrected tilt series. The tilt series were all single-axis from -70 to +50 degrees with approximately two degrees between each projection. The tilt series was reconstructed using the ewbp function included in PRIISM 4.6.1. The reconstructions were binned by a factor of 2 relative to the original tilt series to decrease computation time.

The commands within PRIISM used to compute the reconstruction were:

```
AppendRes tfile 2 1

appl_prm tfile ifile -iprmfile=afile -dimxy=xdim:ydim -iv=0:ntilt:1
    -iref=-1 -imform=2 -pcbase=0.05 -tilt_offset=0 -statfile=none -res=1
    -fullsize=3838:3710 -rscale=2

ewbp ifile rfile -reconxz=1960:960 -sizexz=2000:960 -iy=0:1599
    -filter=2 -hdfilt=1 -moderec=2 -rscale=2
```

where `tfile` is the name of the tilt series file in MRC format and tilt angles in the extended header. `afile` is an alignment file that in these studies specified no further changes to the data. That file is text with two empty lines followed by one line per projection in the tilt series. The $j^th$ of those lines has ten values separated by one or more spaces. The first is the value of $j$. The next eight values are zero, one, zero, zero, one, one, one, and zero, respectively. Those specify no change to the rotation, no change to the magnification, no changes to the translation, and no transformation of the intensities. The final value on the line is the tilt angle, in degrees. `ifile` is the name to use for an intermediate file, in MRC format, that has the tilt series reinterpolated to account for the change in the region of interest for the reconstruction. `rfile` is the name of the file, in MRC format, which will store the weighted backprojection.

**Deconvolving the Weighted Backprojections.** The deconvolution used Muthuvel Arigovindan's ER-Decon II algorithm as implemented in PRIISM 4.6.1. Before the deconvolution, the contrast of the weighted backprojection was reversed, such that densities appeared as bright signal. That was done by multiplying the values by minus one. Since the deconvolution has a background determination step, the value of the constant added when reversing the contrast will only shift the value of the background determined during the deconvolution and not affect the deconvolved volume. The deconvolution was implemented as follows where `ewbp_rev.mrc` is the name of the file with the contrast-reversed weighted backprojection and `tf.mrc` is the name of the file with the normalized transfer function:

```
core2_decon ewbp_rev.mrc deconvolved.mrc tf.mrc -alpha=1e5
    -lamf=5e2 -lampc=0 -lampos=1 -ncycl=100 -linesearch=2014
    -regtype=ma -nonorm_otf -nzpad=zdim -oplotfile=cost_history.txt
    -rfactor
```

The options, `-alpha` and `-lamf`, set the key tunable parameters for the algorithm, $1/\varepsilon$ and $\lambda$ , respectively, as they are denoted in (2). The option, `-ncycl`, sets the number of deconvolution iterations to perform. The `-lampc, -lampos, -linesearch, -regtype`, and `-nonormotf` options were kept constant. The first of those disables the cone filter that the ER-Decon II implementation uses to precondition the problem. The calculations for the cone filter assume that the deconvolution is for optical microscopy and don't match conditions when deconvolving a weighted backprojection. The `-lampos` option specifies the relative weighting for the positivity term in the cost function for the deconvolution. The value used here includes the positivity term with no additional weighting factor in addition to that set by `-lamf` which weights all the regularization terms in the cost function. The `-regtype` option selects the same form for the cost function as in (2). The `-linesearch` option selects a revised line search algorithm. To use the line search algorithm in (2), one would use `-linesearch=2013` rather than `-linesearch=2014`. The `-nzpad` option sets the padded size in z to use during the deconvolution. The value used matches the input size in z so no padding is included. The `-oplotfile` option causes an extra output file to be generated that records the value of the cost function for the deconvolution at each iteration. The `-rfactor` option adds additional data to that file to record the r-factor measure for the blurred deconvolved guess and the input volume to be deconvolved.

**Incorporating a Spatial Constraint in the Deconvolution.** While not applied in this study, a spatial constraint may also be incorporated in the deconvolution, implemented by adding a term to the cost function that is the sum over positions of the squared product of the current guess, g(r), with a spatial mask, M(r),

$$\lambda\lambda\_w\Sigma\_r(g(r)M(r))^2$$

Where the mask is not zero, nonzero intensities in the guess are penalized. The first scaling factor multiplying the sum is the overall scaling factor, $\lambda$, for the regularization terms added to the cost function. The second scaling factor, $\lambda$w, controls the relative weight of the spatial constraint to the other regularization terms. Setting that scale factor to zero nullifies the spatial constraint so the cost function is identical to the one in (2). The addition of the term to the cost function changes equation S22 in (2) paper by adding a term,

$$\lambda\lambda\_wg(r)M(r)$$

, to the left hand side. It also changes the diagonal approximation in equation S29 by adding a term,

$$\lambda\lambda\_wg(r)M(r)$$

, to the right hand side. For the computation of the initial guess, contribution from the spatial constraint is ignored.

An implementation of the ER-Decon II algorithm with the spatial constraint term is available in PRIISM 4.7.0.

For the experiments performed, the mask for the spatial constraint was not changed during the deconvolution iterations, and a value of one was used for λw. The command options to include the spatial constraint in PRIISM's `core2_decon` are:

$$-lammask = 1 - imask = mask\_file$$

where `mask_file` is the name of the file, in MRC format, that contains the values for the spatial constraint mask. The mask must match the dimensions of the padded input volume.

**Constructing a Spatial Constraint Mask.** A hard binary mask was constructed for the spatial constraint as follows:

1. In every 5th slice perpendicular to the tilt axis of the deconvolved weighted backprojection, image coordinates are selected corresponding to the top and bottom edges of the slab of material appeared to intersect the left and right edges of the deconvolved weighted backprojection.

2. A linear least squares fit is found for each of the four sets of coordinates for the corners of the slab, upper left, upper right, lower right, and lower left to define the boundaries of the volume.

3. For each slice perpendicular to the tilt axis the vertices of the parallelogram whose left and right sides match the left and right edges of the deconvolved backprojection are calculated using the linear models from step two for the coordinates of each corner.

4. For the spatial constraint mask in a slice perpendicular to the tilt axis, any sample outside the parallelogram computed in step three, is set to one, and any sample inside the parallelogram is set to zero.

**Perturbing Deconvolution Iterations.** Some experiments stopped the deconvolution after a certain number of cycles, modified the guess, and then used the modified guess as the starting point for more deconvolution iterations. The implementation of ER-Decon II in 4.7.1 allows the initial guess for the deconvolution to be provided as a file in MRC format. The command line option that sets the initial guess is

```
-guess=file_name
```

where `file_name` is the path to the file, in MRC format, with the values for the initial guess.

**Computing Fourier Transform Results.** Fourier transform results for the undeconvolved or deconvolved weighted backprojection were computed as follows:

1. A 400 x 400 x 400 box was extracted whose bottom lower left corner was at the sample coordinates, (300, 40, 200), where the first coordinate is the direction perpendicular to the tilt axis and axial direction, the second coordinate is in the axial direction, and the third coordinate is along tilt axis.

2. The mean of the extracted volume was subtracted from that volume's values.

3. The result of step two was apodized by applying a triangular window to each dimension. The triangular window is zero at the first sample in the dimension, linearly increases to one at the 200th sample, and then linearly decreasing to .005 at the last sample.

4. Three-dimensional discrete Fourier transform of the result of step three was computed and, for display purposes, the zero frequency term was shifted to be at sample coordinates (199, 199, 199) in the 400 x 400 x 400 volume.

**Computing the Cylindrically Averaged Fourier Transform Amplitudes.** The procedure to compute the cylindrical average, an $nr \times nz$ element array called `cyla`, of the Fourier transform amplitudes in a $n \times n \times nz$ array was as follows:

1. Set $nr$ to be $floor((n-1)/2)$, where $floor(v)$ is the largest integer less than or equal to $v$.

2. For each plane, $iz$, in the transform perpendicular to the axial frequency axis:

   (a) Initialized a $nr$ element counter array, $icount$, to zero and a $nr$ element amplitude sum array, $asum$, to zero.

   (b) For each element, coordinates (ix, iy), in the plane:

    i. Computed the amplitude, $a$, of the Fourier transform.
    ii. Compute the radial frequency, $k$, from the coordinates. Since the zero frequency component in $x$ and $y$ had coordinates of $(nr-1, nr-1)$ and the pixel spacing and dimensions in $x$ and $y$ are isotropic, $k$ is $\sqrt{(ix-nr+1)^2+(iy-nr+1)^2}$.
    iii. If $k$ is less than $nr$ , let $ik$ be $floor(k)$. Add one to $icount[ik]$ and add to $asum[ik]$

3. Set $cyla[iz, j] = asum[j]/max(icount[j], 1)$ for $j$ between 0 and $nr-1$, inclusive.

For display in the figure, only every other frequency, from the zero frequency, was displayed.

Using commands from PRIISM, the computation of the Fourier transform of a 400 x 400 x 400 volume from v.mrc, which has the full weighted backprojection or deconvolution of the weighted backprojection, and computation of the cylindrical average, ad.mrc, of the amplitudes was:

```
FTransform3D v.mrc f.mrc -x=300:699 -y=40:439 -z=200:599 -subtract=mean
    -triangular=0.5:0.5:0.5  -real_complex_full
    -center_zero -same_units
Flip f.mrc ff.mrc -xz
RadProj ff.mrc af.mrc -format=mrc -polar -average -components=amplitude
    -center=199:199:199 -r=0:200 -nr=200 -theta=-180.01:180.01 -ntheta=1
Flip af.mrc a.mrc -xz
Decimate a.mrc ad.mrc -x=0:199 -y=1:399 -factor=2:2
```

`Flip` was used after the Fourier transfer because the output of the weighted backprojection has the axial dimension as the second fastest varying axis in the file. For the averaging step, it needs to be the third fastest varying axis. The second use of `Flip` wass to convert a nr x 1 x nz file to a nr x nz x 1 file.

### Notes on the DC Methods

If the centering convention for the tilt series alignment is not consistent with the conventions used by `appl_prm` and `ewbp`, that would introduce some error in the reconstruction. `ewbp` in version 4.7.1 of PRIISM has ways to override its default centering conventions so one could correct for a mismatch without regenerating the aligned projections.

The placement of the projection of the point object in the synthetic projections used to compute the PSF does not exactly match up with `ewbp`'s centering conventions. By default, `ewbp` places the x coordinate of the center of the aligned projections at 0.5 * (nx_projection – 1) and assumes the projection rays through the centers intersect at (0.5 * (nx_recon – 1), 0.5 * (nz_recon – 1)) in the xy planes of the reconstruction. Since the projection and reconstruction dimensions used here are even, those conventions put the x coordinate of the center of the projection at 499.5 and the center of the reconstruction in the xz plane at (499.5, 239.5). For the synthetic projections, the center of is defined at 499.

A separate deconvolution was performed using `ewbp` 4.7.1 to override the default centering convention. The deconvolution result from that transfer function was, qualitatively, much the same as the transfer function generated with the procedure in the methods. For future work, `ewbs`'s conventions in the computation of the transfer function should be overridden to avoid the mismatch. That would mean replacing:

```
ewbp synproj_blur.mrc psf.mrc -reconxz=1000:480 -sizexz=1000:480 -iy=0:799
    -filter=2 -hdfilt=1 -moderec=2
```

with:

```
ewbp synproj_blur.mrc psf.mrc -reconxz=1000:480 -sizexz=1000:480 -iy=0:799
    -filter=2 -hdfilt=1 -moderec=2 -pcen=499 -reconcen=499:399
```

The shift specified for the Fourier transform of the PSF would also change. Replace:

```
FTransform3D psf.mrc tf.mrc -same_units -shift=499:240:399
```

with:

```
FTransform3D psf.mrc tf.mrc -same_units -shift=499:239:399
```

The computation of the PSF assumes that there's something in the system that will guarantee that the point object ends up being band-limited to the frequency range set by the pixel spacing. If one isn't willing to assume that, one could generate synthetic projections that have 1/m times the pixel spacing and have m times the samples in each dimension, apply the microscope CTF as desired, reconstruct the synthetic projections, and then finally bin the result by m in each dimension. For a sufficiently large m, likely 3 or 4 in this case, that would account for the high frequency information that would be aliased into the measurement.

Once the PSF has been generated, a range of DC parameters are tested and compared to choose which combination of smoothing and non-linearity parameters is optimal (SI Appendix Fig S2, S S3, our choice for optimal combination highlighted in blue). The optimal parameter combination is judged based on the filling of the missing wedge as seen in Fourier space, as well as the change in contrast in the real space image. We typically find the optimal smoothing parameter to be between 400-600, and the non-linearity parameters give optimal results around 10000.

### New DC Directions

There is no question that the DC process changes intensity distributions, sharpening features and minimizing fog/noise. The process does not alter the monotonic relationship of the intensities (i.e. does not invert contrast), only the relative difference in intensity. Flips of intensities from positive to negative or the reverse have never yet been observed in the DC process. Still, intensity changes must be interpreted with caution. Do all areas of the intensity histogram contain similar amounts of information post DC? For example as a function of scaling the DC image and comparing the bright areas of the histogram to weak intensity areas, are there differences in Fourier space DC representation ( see especially (3))? There are additional aspects of the DC process, such as extension of the iteration number( up to 1000 iterations), and other DC steps that will be addressed in future work.

It is important to carefully scrutinize DC images for differences, artifacts, or errors, as this procedure is still new. One does not know if the parameters for the cytoplasm are different from those optimal for the nucleus, for example. One does know that once parameters are known for a region of the tomogram, it is possible to move the DC area/volume around the large tomogram and get equivalent DC. Still, the optimal parameters have been similar for all tomograms deconvolved.

Could the DC process reduce the electron dose required for the cryo-ET? With the improvements gained from DC, it is conceivable that the number of tilt steps required might be reduced. Alternately, a lower electron dose could potentially be used, followed by image restoration by DC. DC could also enable the study of slightly thicker volumes (200-400 nm) by reducing the noise inherent in these thicker images. It is known that for this thickness range, about half of electron scattering events are inelastic. These inelastically scattered electrons are separated from the elastic (zero loss) scatter events by an energy filter at great effort/ expense. The inelastic events do not contribute to usable image information, creating instead a fog of background noise. Might the DC improve tomograms obtained without an energy filter?

The DC process might also benefit from a large reduction of defocus, while still recovering the contrast. The PSF of this lower defocus tilt series would be much simpler. If this were possible, DC could act as a computational substitute for the expensive/complex phase plates.

For computing how the defocus varies across an image with a nonzero tilt angle, $pfocusramp$ assumes that the defocus parameter you supply correspond to the defocus near the center of the image (zero-based pixel coordinates, $(floor(nx/2) - 1, floor(ny/2) - 1)$, to be precise). At the pixel, $(ix, iy)$, in zero-based indices, $pfocusramp$ will add to the defocus amount.

```
p * sin(t) * dot_product(
  (ix - floor(nx/2) - 1, iy - floor(ny/2) - 1),
  (cos(a), sin(a))
)
```

where a is the tilt axis orientation angle, t is the tilt angle, and $p$ is the pixel spacing. In other words, if the tilt axis is vertical $(a = 0)$, the tilt angle is positive, and the center of the image is underfocused, the points to the right of center will be less underfocused; points to the left of center will be more underfocused.

Lastly, while the DC process is computationally reasonable time frame, a rewrite into modern GPU technology would greatly speed up the computational time.

### Movies, and Their Discussion

Because the advantage of DC relates to improvements in 3-D resolution, it is important to visualize and interpret the data in 3-D. One effective way to visualize DC data is to use stereo pairs (SP) where two angular views are displayed side by side, and viewed such that the brain interprets the two images as one 3-dimensional scene. To do this, a 3-D region (cube) of interest is selected from the original 3-dimensional DC image. Two projections of that cube are then generated, with a 6-12 degree difference in the projection angle. Then, by viewing the SP straight on (12-24 inches away) with slightly crossed eyes, one can visualize a third image of the volume in 3D.

Viewing a volume in motion can also aid in interpretation of structures in the DC volume. By animating the rocking of an SP (rotating angle stereo pair, or RASP) while visualizing the cube in 3-dimensions, the SP are rotated in the computer and the SP, offset in 6-12 degrees, are generated and viewed. In addition, the cube of data can be tilted at several large angles to generate multiple angular views, each in different orientations, rocked and in stereo (SP). We recommend to generate and visualize movies in stereo, and take advantage of the play bar to allow control of the rocking. Information in the SP does require careful study(and considerable time to perceive the structures). The DC process is not perfect, and the differences in resolution (in Z, for example) help many times to reconcile and bring into perspective the different structures in the 3-dimensional volume.

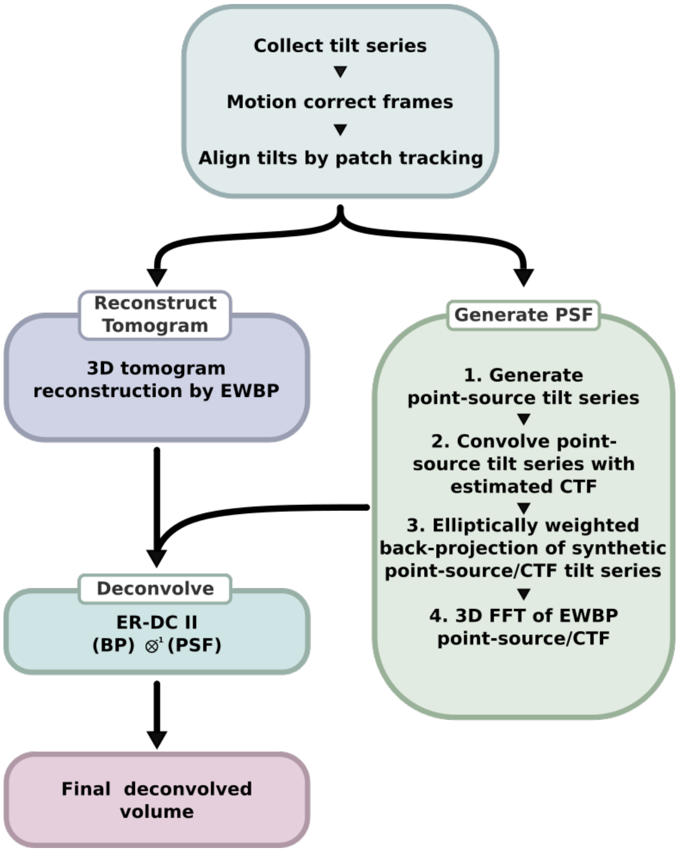


**Fig. S1.** Schematic of the deconvolution workflow.

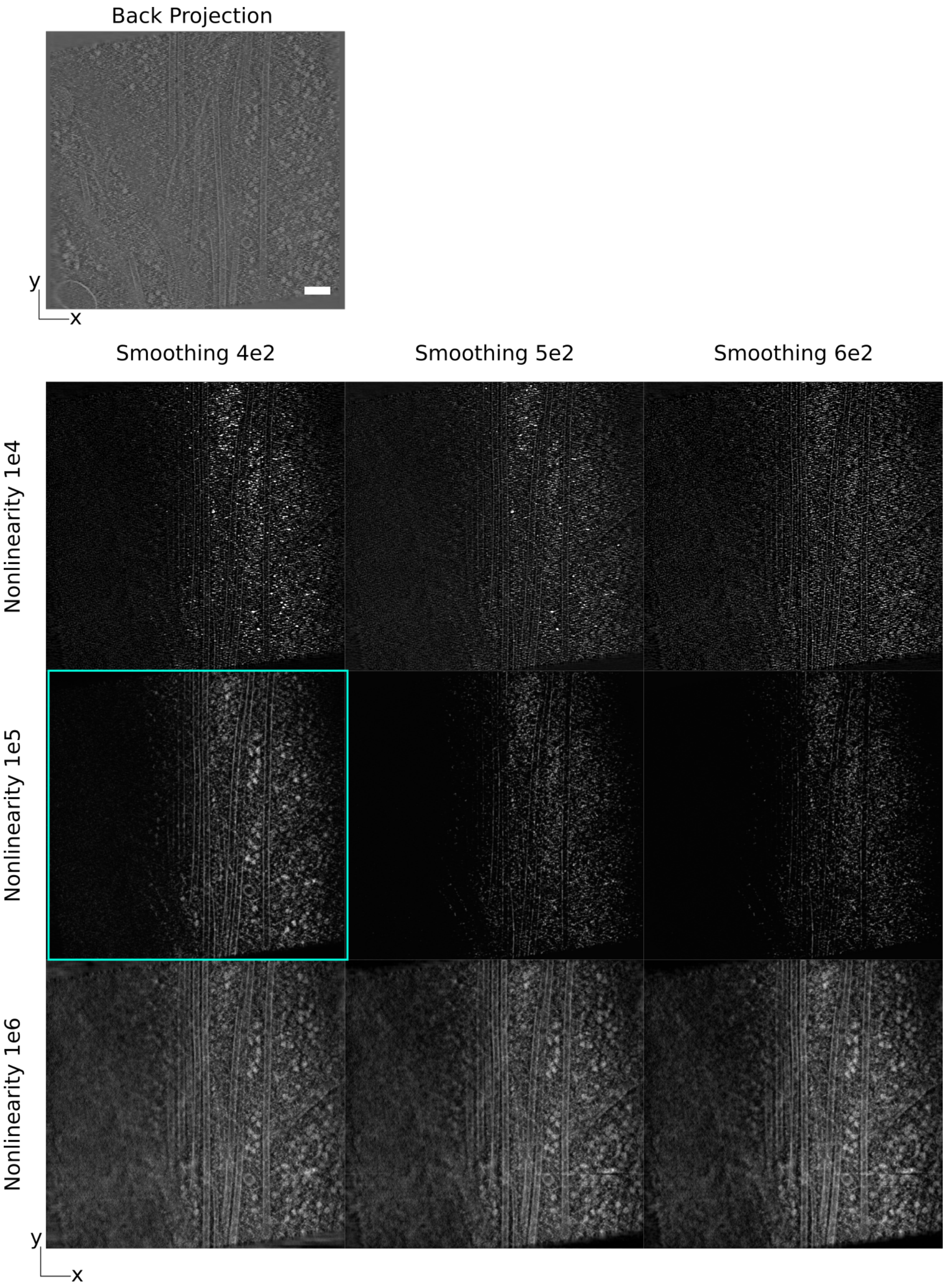


**Fig. S2.** Matrix of deconvolution results at different smoothing and non-linearity parameters (XY projections). Scale bar: 100 nm.

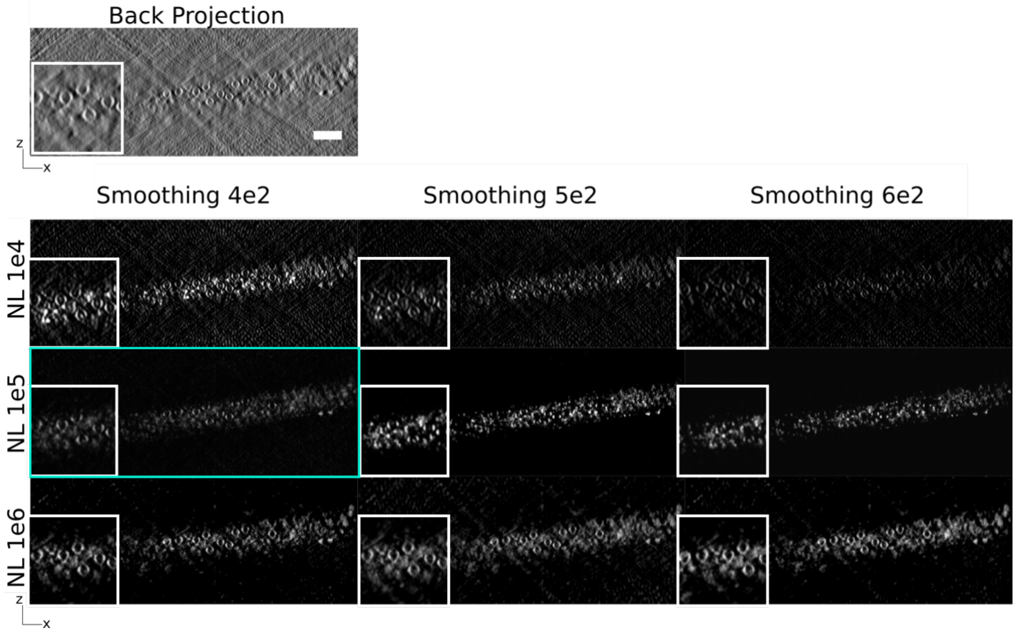


**Fig. S3.** Matrix of deconvolution results at different smoothing and non-linearity parameters (XZ projections).Scale bar: 100 nm.

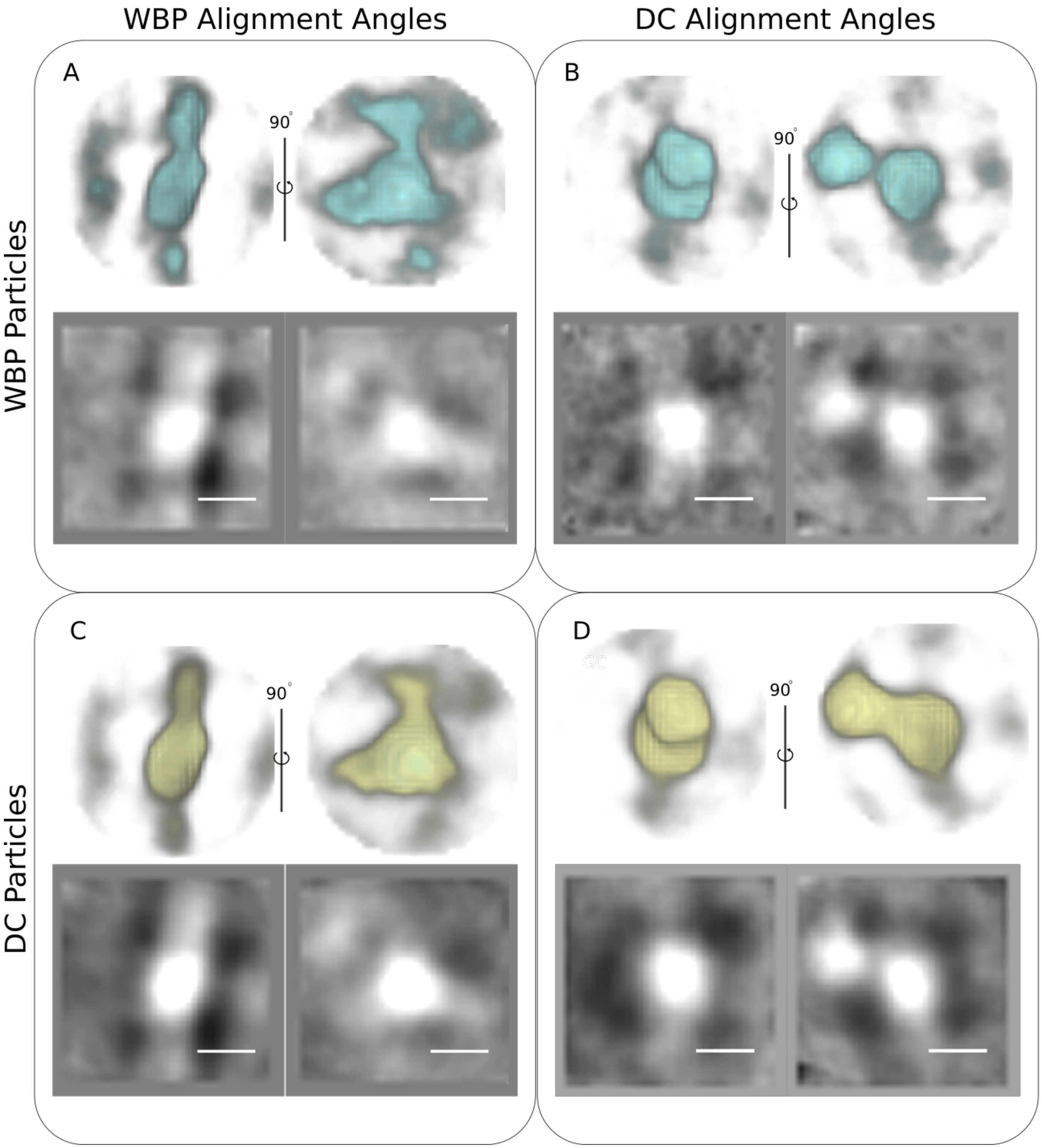


**Fig. S4.** Comparison of WBP aligned particles and DC aligned particles. (A) Volume views of the crystal body average generated by aligning WBP subtomograms. (B) Volume views of the crystal body average generated by aligning WBP subtomograms using alignment parameters generated by aligning DC particles. (C) Volume views of the crystal body average generated by aligning DC subtomograms. (D) Volume views of the crystal body average generated by aligning DC subtomograms using alignment parameters generated by aligning WBP particles. Scale bar: 10 nm

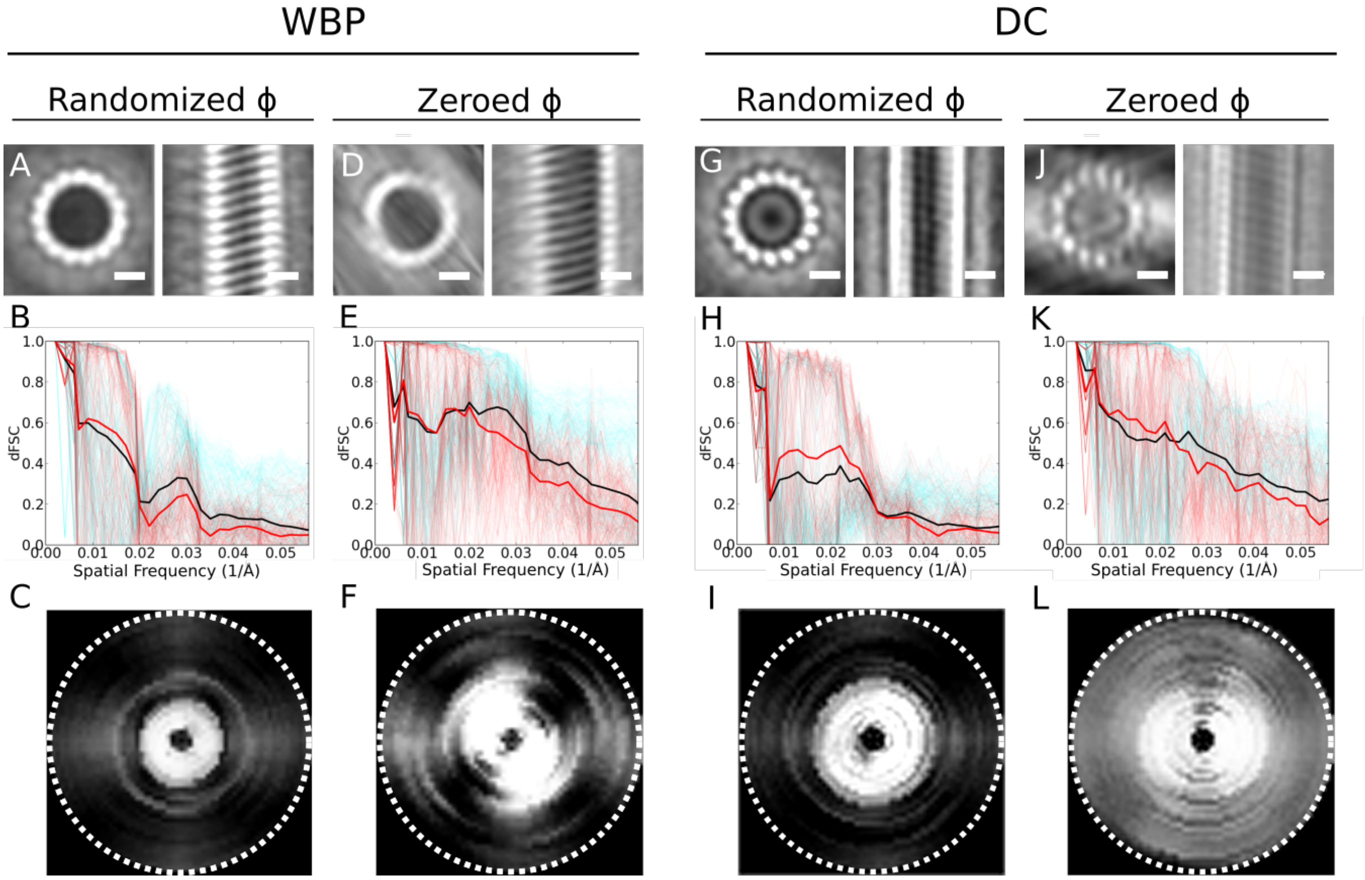


**Fig. S5.** Microtubule subtomogram averages and their dFSCs.(A) Average from the WBP tomogram, using a randomized azimuth-angle ($\phi$) averaging approach to compensate for the missing wedge (4). (B)dFSC corresponding to (A). (C) Central slice of the dFSC from (B). (D) Average from the WBP tomogram, with initial constant $\phi$ angles for all particles, allowing the missing wedge to drive the alignment. (E) dFSC corresponding to (D). (F) Central slice of the dFSC from (D). (G) Average from the DC tomogram, using a randomized azimuth-angle ($\phi$) averaging approach to compensate for the missing wedge. (H) dFSC corresponding to (I). (I) Central slice of the dFSC from (H). (J) Average from the DC tomogram with initial constant $\phi$ angles for all particles, allowing the missing wedge to drive the alignment.(K) dFSC corresponding to (J). (L) Central slice of the dFSC from (K). Cyan lines in plots represent dFSCs of angles sampled during tomography. Red lines in dFSC plots represent those angles that comprise the missing wedge, thick red line is their average. Black line represents the global average of all dFSCs. Scale bars: 10 nm. Dashed circle indicates Nyquist resolution, 0.059$\text{Å}^{-1}$.

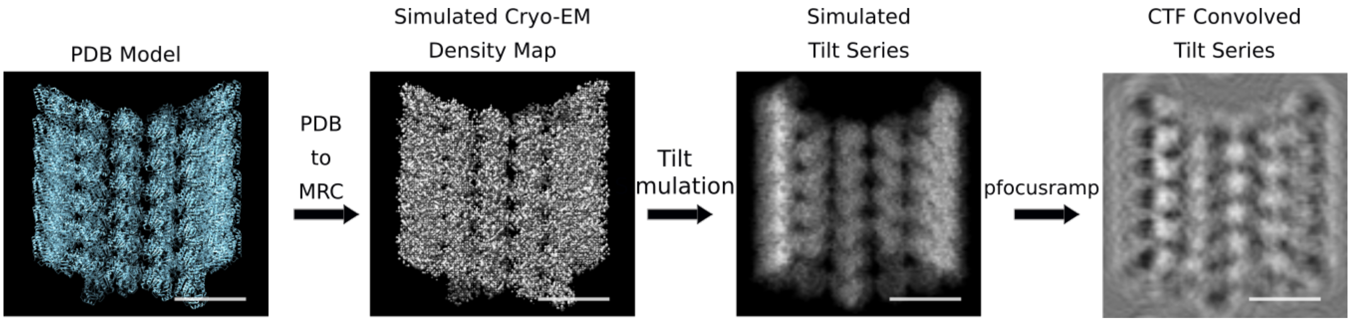


**Fig. S6.** Workflow for simulation of tilt-series. A simulated cryo-EM density (MRC format) is created from an atomic model of a microtubule (PDB format). Then, a tilt series of is simulated from the map and then convolved with a -3.00 $\mu$m defocus CTF. Scale bar: 10 nm.

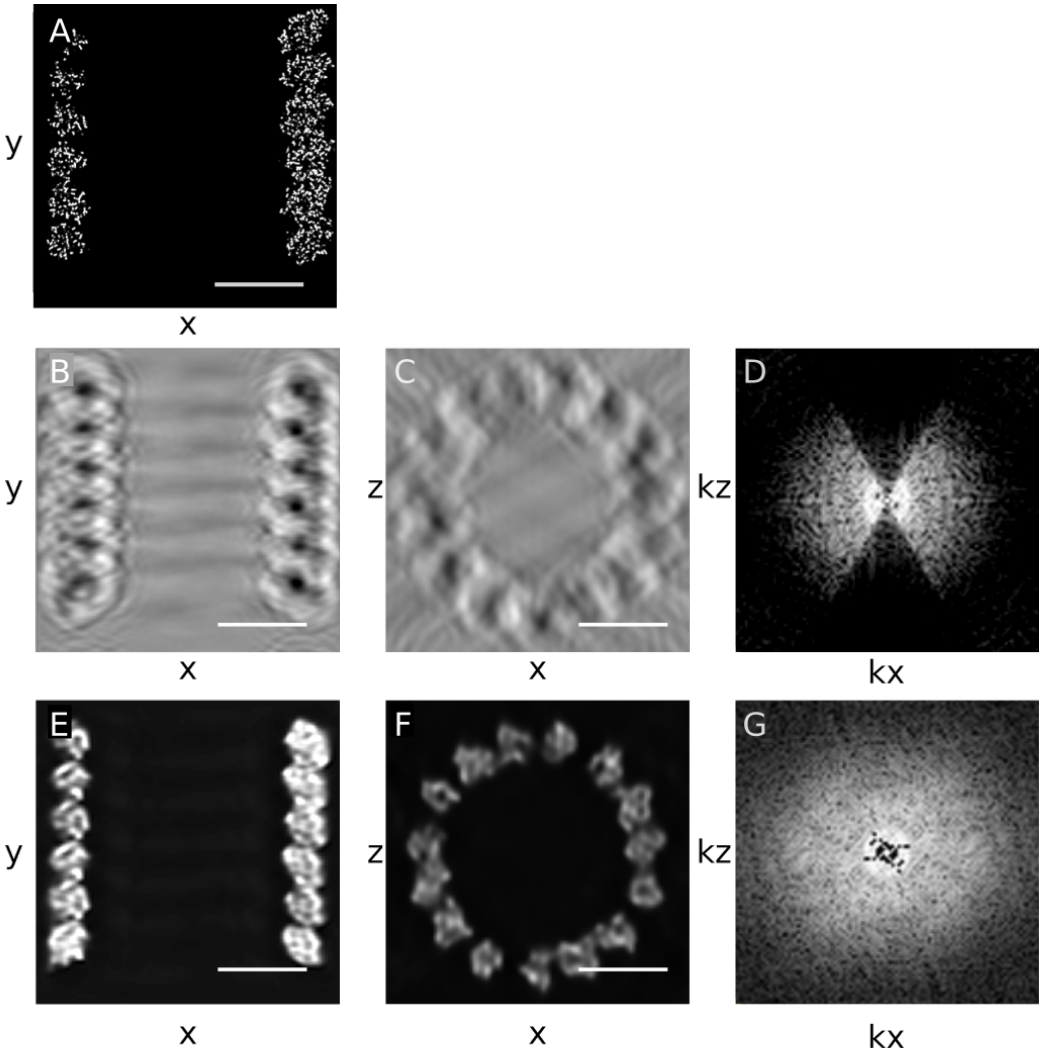


**Fig. S7.** Back projection of simulated tilt series and corresponding deconvolution. (A) Central slice of a simulated the microtubule cryo-EM map. (B) XY central slice of the tomogram reconstructed using WBP from the CTF-convolved simulated tilt series. (C) XZ central slice of the tomogram in B. (D) kXkZ fourier spectrum of (C). (E) XY central slice of the deconvolution. (F) XZ central slice of the deconvolution. (G) kXkZ fourier spectrum of (F). Central slice thicknesses are all 10 Å. Scale bar: 10 nm

**Movie S1. RASP1. Rotating stereo-pair of crystal body WBP subtomogram at different angles.**

**Movie S2. RASP2. Rotating stereo-pair of crystal body DC subtomogram at different angles.**

**Movie S3. RASP3. Rotating stereo-pair of yeast WBP subtomogram at different angles.**

**Movie S4. RASP4. Rotating stereo-pair of yeast DC subtomogram at different angles.**
